# Automated benchmarking with structured variant effect data

**DOI:** 10.64898/2026.11.27.769673

**Authors:** Floris J. van der Flier, Ethan Eschbach, Karel van der Weg, Thomas Hopf, Cor Zuurmond, Shu Zhao, Prashant Srivastava, Sergio Garcia Busto, Charles W. J. Pugh, Jonathan Frazer, Mafalda Dias, Daniel Probst, Dick de Ridder, Pascal Notin, Debora S. Marks, Henning Redestig

## Abstract

Protein variant effect prediction (VEP) has a wide range of applications, from describing the role of mutations in human disease to improving enzymes for industrial applications. VEP is now predominantly powered by machine learning models and high-throughput experimentation. Progress in this field depends on fair and systematic model comparison, which is complicated by the heterogeneity of VEP tasks and the diversity of model implementations. ProteinGym is the most widely used VEP benchmark, but it does not have a standard representation for datasets or models. This makes it difficult for others to run their own benchmarks with it and for the benchmark to keep up with new datasets and models. We address these shortcomings with the ProteinGym framework (PGF), which standardizes dataset representations, model interfaces, and evaluation protocols. The PGF consists of two packages: `proteingym-base`, which provides schemas for composing datasets and the PGDATA archive format, a standardized representation of VEP datasets, and `proteingym-benchmark`, which provides automated benchmarking of containerized models. As an example of a new experiment, we use the PGF to measure the ability of three supervised models to identify variants with property values that exceed those in the training set, a common objective in protein engineering. The PGF is available at https://github.com/ProteinGym.

## 1 Introduction

The prediction of protein variant effects (VEP) is essential for a range of applications within the life sciences, including the description of the role of mutations in disease and the identification of improved enzyme variants for industrial processes [1, 2]. Models are developed in two settings; zero-shot predictors score variants by learning constraints from evolutionary, structural, or functional data [3, 4, 5], while supervised predictors are trained on measurements from an assay of interest and evaluated on held-out variants of a similar protein [6, 7]. In both settings, the community continues to develop methods that use new combinations of sequence, structure, and alignment information [8, 9, 10]. Model comparison is essential both to guide the development of new models and to select the most suitable one for a given VEP application; yet it remains laborious. Implementations are diverse and developers often benchmark their models on different datasets with different scoring procedures. In applications, scientists may prioritize convenience over performance. When a prediction task is well defined, different models can be compared and improved iteratively, as demonstrated by CASP for structure prediction [11] and CAGI for genome interpretation [12].

For supervised protein sequence-function modeling, benchmarks such as TAPE, FLIP, and PEER define tasks with fixed data splits [13, 14, 15], and deep mutational scanning data have been used to benchmark variant effect predictors at scale [16, 17]. ProteinGym was created to address the issues that arise from the lack of formalization in VEP [18]. It encompasses a large number of phylogenetically and phenotypically diverse protein function assays, as well as a large set of human genes with clinical annotations, both curated with extensive metadata. More than 70 VEP models were initially benchmarked across all datasets. Zero-shot predictors were scored on the full assays, and supervised predictors were evaluated by cross-validation (CV), for which “modulo” and “contiguous” schemes were introduced alongside random CV to probe each model’s ability to extrapolate in the sequence domain. The results were organized on a public leaderboard that allows analysis of the performance of the model by the assay function, taxon, and alignment depth. This allowed developers to benchmark their models as they were released [7, 19, 20] and researchers to analyze the benchmarking results for dataset and model characteristics of interest [21].

Two gaps in the implementation of the original ProteinGym complicate its adoption. First, ProteinGym does not impose a data standard. Models each expect their own data format, which means that every model needs a bespoke adapter script maintained inside the ProteinGym codebase. Model builders have no specification to follow, and datasets are not validated for correctness. Second, the pipeline runs the full matrix of models against datasets within a single execution environment and cannot easily be filtered down to a subset of either. In addition, the leaderboard ranks models by their average scores without testing whether the differences between them are significant. In our view, ProteinGym in its current form is better suited to reporting benchmark results than to the iterative refinement of models or model comparison.

Addressing the first gap calls for standardized models and datasets. VEP datasets are distributed as CSV files that are decoupled from their metadata, and existing efforts to add structure do not deliver a self-contained representation for machine learning. EnzymeML offers schema-based validation and packaging of enzymology experiments [22], but focuses on documenting the experiments rather than packaging the data for model training. MaveDB provides curated variant effect data without bundling the artifacts (structures, MSAs, sequences) that models require [23]. Models are similarly unconstrained, and without a shared model interface published models generally cannot be integrated into a workflow without manual intervention. The second gap calls for tooling that runs each model in its own isolated environment. Without isolation, a pipeline becomes less maintainable with every added model, as dependencies accumulate and conflict, and individual jobs cannot be dispatched to dedicated hardware.

To address these gaps, we introduce the ProteinGym framework (PGF), an open-source software suite for automated benchmarking of VEP models. Its two design principles are standardized datasets and model interfaces, and containerized model execution. The PGF comprises (i) `proteingym-base`, which defines the PGDATA archive format for datasets together with tools to split and score them, and (ii) `proteingym-benchmark`, which runs parameterized benchmarking pipelines, validates contributed models through continuous integration, and publishes results to a web catalog that tests whether differences between models are significant. The PGF can take advantage of the evedesign model interface, a framework for biosequence design [24], so that every benchmarked model is driven by the same entry point and can be used directly for design purposes. The initial release contains 12 zero-shot and 4 supervised models. To show how the framework can be extended, we tested the ability of three supervised models to identify variants with property values that exceed those of the training data.

## 2 The ProteinGym Framework

### 2.1 The ProteinGym framework separates datasets, models, and benchmarking

The PGF has two layers (Figure 1). The first, `proteingym-base`, defines an archive format for VEP datasets, tools to split and score them, and a model card that describes how a model is configured. The second, `proteingym-benchmark`, runs containerized models against datasets in automated pipelines, validates newly contributed models, and publishes the results. Datasets are distributed through a Hugging Face Datasets repository [25]. Datasets, models, and results are presented in a web catalog. All models and datasets satisfy the same contracts; a new model enters the public benchmark through a pull request, and a new dataset enters it once it is packaged with `proteingym-base`. As illustrated in Table 1, PGF is unique in the sense that containerisation and contracts facilitate maintenance and local benchmarking experiments.

**Table 1:**
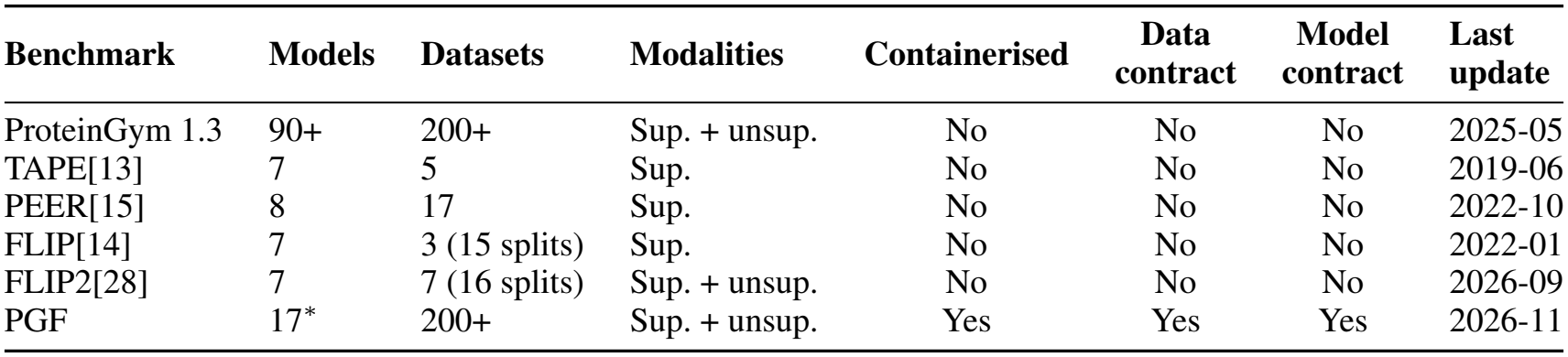
Comparison of PGF with other VEP benchmarking tools. PGF’s containerisation and its model and data contracts make it easier to maintain and to run locally. Sup. = supervised; unsup. = unsupervised (zero-shot).

**Figure 1:**
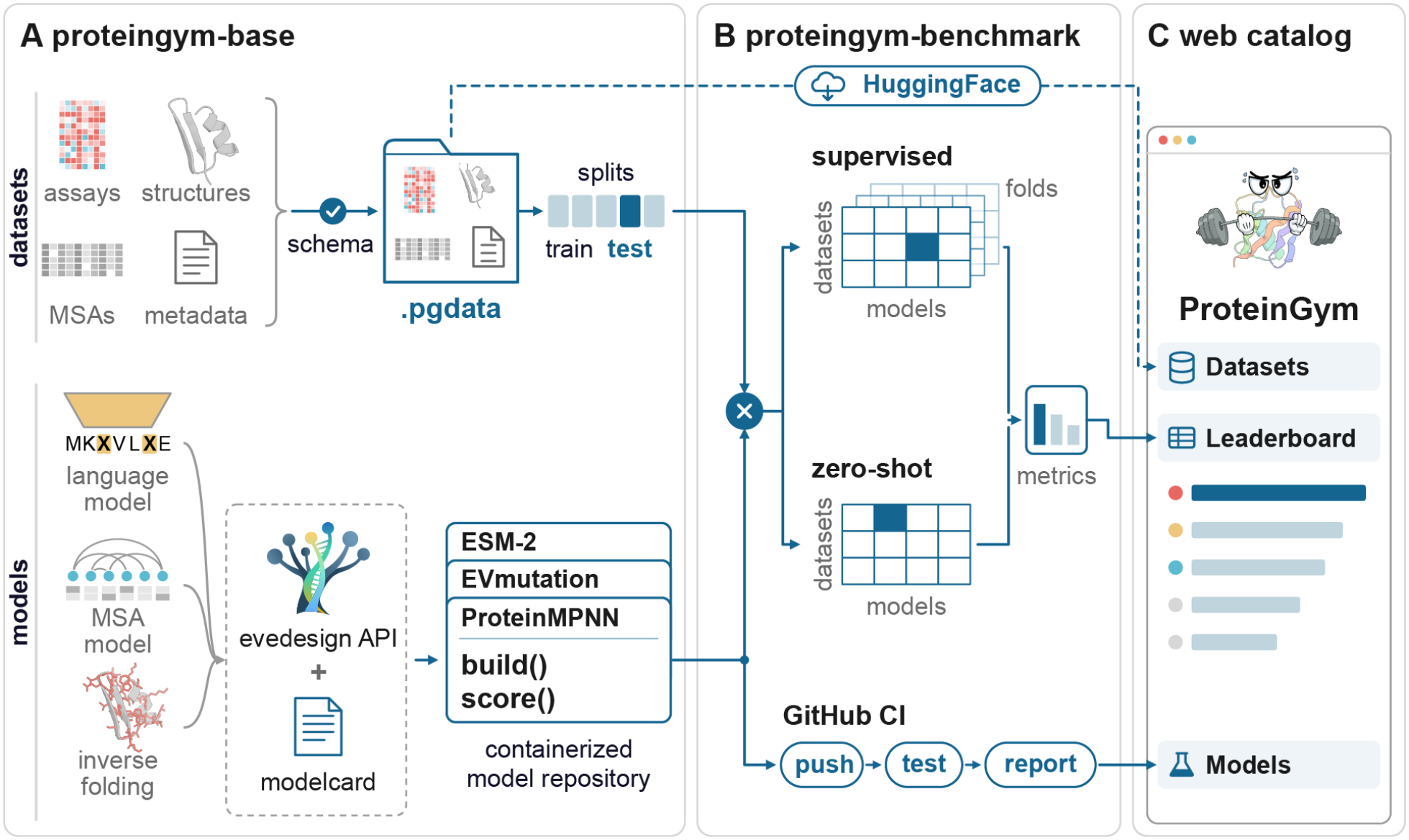
The ProteinGym framework comprises two components. (A) The `proteingym-base` repository provides an installable package for creating standardized datasets validated by schemas and for describing models with a model card. Datasets come in the form of PGDATA files, which are archives containing assay data, structures, MSAs, and metadata, and can be partitioned into train and test splits. Models are optionally wrapped with the evedesign API, which exposes standard `build` and `score` operations, and are distributed as containerized model projects. (B) `proteingym-benchmark` runs benchmarking experiments over all combinations of models, datasets, and folds, separately for the supervised and zero-shot settings, and validates contributed models with a GitHub CI workflow. Datasets in PGDATA format are hosted in a Hugging Face Datasets repository. (C) Published models, datasets, and benchmarking results are displayed in a web catalog.

### 2.2 PGDATA packages VEP datasets into validated, self-contained archives

VEP models are trained on different data modalities, including assay measurements, multiple sequence alignments (MSAs), and 3D structures, each of which is stored in a variety of specialized formats (e.g., CSV, A2M, and mmCIF/PDB, respectively). These files are usually distributed separately from the metadata that describes them and are not validated against each other, so training or testing a model on a new dataset typically requires bespoke reformatting. With `proteingym-base`, we package all modalities of a dataset, together with its metadata, into a single validated archive in the PGDATA format.

Each dataset is defined by a human-readable `manifest.toml` file written by the dataset curator. The manifest documents the dataset, including relevant publications, the measured properties (assay targets) and their units, and the experimental conditions (assay variables) of each assay. The manifest also serves as a recipe that lists the files from which `proteingym-base` constructs the archive. It follows a versioned schema of mandatory and optional components; every data set must contain at least one assay, while structures, MSAs and raw assay files are optional (Supplementary Table S1, Supplementary Listing S1). During construction, we parse each component into Pydantic [26] data classes for validation. This catches errors such as missing target columns or incompatible sequence alphabets at the time of dataset creation, before any model is run.

The resulting PGDATA file is a zip archive with a fixed layout. It contains one or more assay tables of variant sequences and target values, which allows a publication reporting several assays to be represented as a single dataset. Next to the assays, the archive holds the artifacts that models may require, such as reference sequences, structures, MSAs, and MSA sequence weights. A machine-generated lock file, a resolved copy of the manifest, records all metadata and the location of every artifact inside the archive. Data splits share the same representation, as splitting strategies partition the assay data into slices that are stored with the dataset as a PGDATA archive. `proteingym-base` currently supports random, *K*-fold, quantile-based [27], and predefined splits, the latter of which preserve the modulo and contiguous splits provided in the original ProteinGym release [18]. Because splits are stored with the data rather than computed by each model, all models are trained and tested on identical partitions.

### 2.3 Models share a single interface and run in isolated containers

The heterogeneity of VEP models hinders benchmarking on two fronts. Technically, models do not share an interface, and their software dependencies may conflict; conceptually, zero-shot and supervised predictions exist on different numeric scales, which complicates interpretation when both are combined. We address the first issue with a shared model interface and containerized execution, and the second by benchmarking the two settings in separate tracks (Section 2.4).

A model interface defines how a model receives data and how it returns predictions. In existing VEP benchmarking projects, every model requires a bespoke adapter between the dataset and its code, and the resulting code, which we refer to as a wrapper, cannot be used outside of the benchmark. We consider it preferable to standardize the operations that models perform, which allows a single entry point to drive all of them. We adopted the interface of evedesign, an open-source framework for biosequence design [24], which frames design as a conditional modeling problem. In this formulation, a model is first built on a molecular system that carries any available information, such as the wild-type sequence, homologous sequences, and structures, optionally together with labeled training data, and the conditioned model then scores instances, i.e. concrete variants of the system. These two operations correspond directly to fitting a model and predicting variants, and apply equally to zero-shot and supervised predictors. We chose evedesign because it has reference implementations of many models relevant to VEP, its representation of systems and instances maps directly onto the contents of a PGDATA archive, and any model wrapped for the benchmark is also available for design workflows. The PGF does not depend on evedesign, and another model framework with equivalent operations could replace it.

All models in the current release are driven by the same entry point (Figure 2A). Each model is described by a model card [29], a `README.md` file that documents the model and carries structured front matter with its name, the class that implements it, a tag declaring its prediction setting, and its hyperparameters (Supplementary Listing S2). Given a PGDATA dataset, an assay prediction target, and, for supervised models, a split and a test fold, the entry point converts the dataset into an evedesign system with training and test instances, instantiates the model class with the hyperparameters from the model card, instantiates the model, and scores the variants (Supplementary Section A.3). Supervised models are trained on the training folds, whereas zero-shot models receive no training data and score the full dataset. Like the model wrapper, the entry point could be replaced by a different implementation of the same model interface.

**Figure 2:**
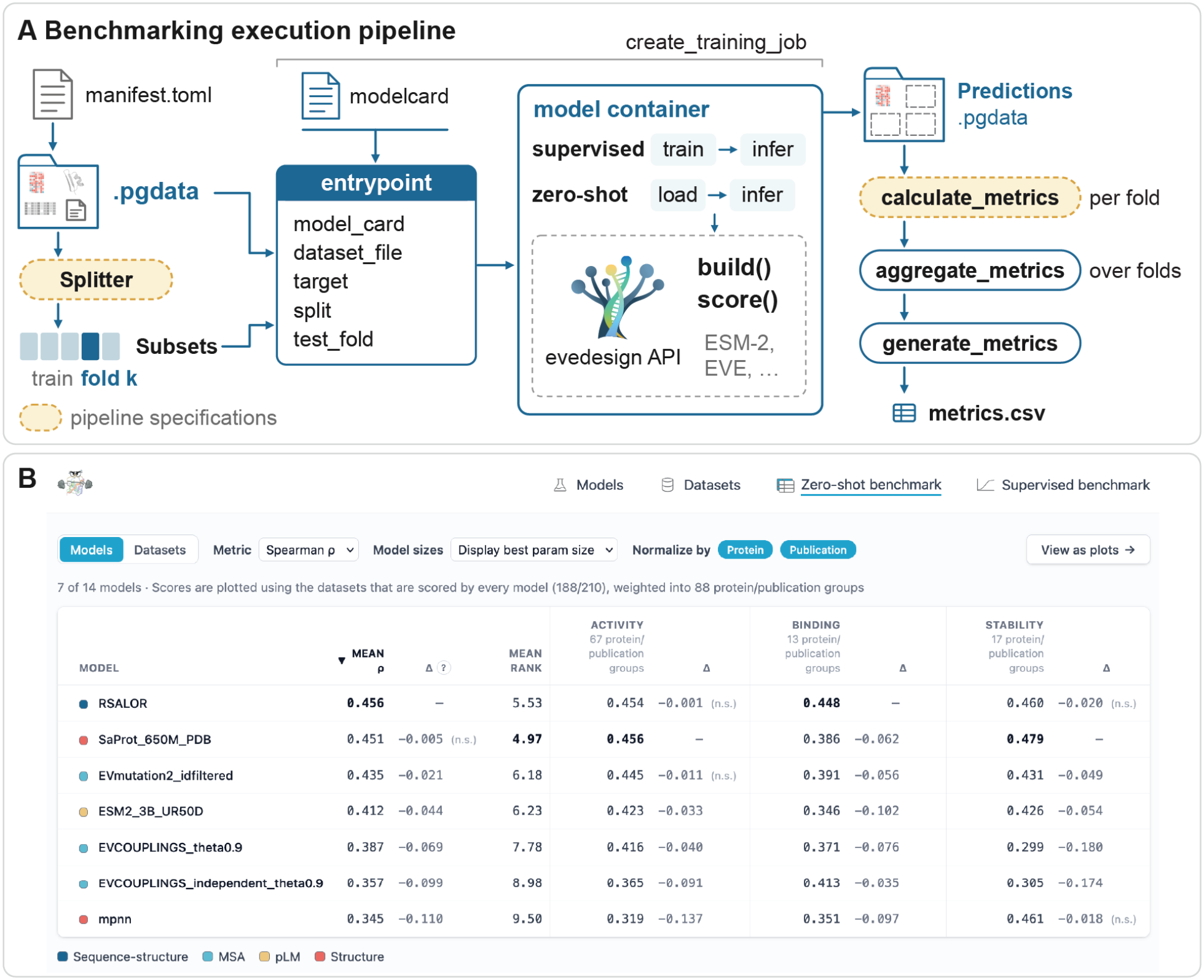
Benchmarking pipelines of the ProteinGym framework. (A) A benchmarking job passes a dataset, its splits, and a model card to a standard entry point, which drives the containerized model through the evedesign API and writes predictions as a PGDATA file. Predictions are scored per fold, aggregated over folds, and summarized in a single table. Dashed boxes indicate components that are specified per experiment. (B) The web catalog benchmark result table displays model results with rigorous statistical comparison.

Models return their predictions as a PGDATA file derived from the input dataset, which preserves the structure and record order of the input assays, replaces the measured values with predictions, and removes all artifacts (MSAs, structures, sequences) and metadata. Preserving record order means that the slices that define the CV folds of the input apply unchanged to the predictions, which allows scoring to take place outside the model container with the same code for every model. Removing the artifacts keeps the output small. Variants that a model cannot score, e.g. insertions and deletions for fixed-length models, are left without a prediction. Predictions are matched with records by sequence. In an unsupervised setting, a sequence measured under several assay conditions currently receives a single prediction for all of them.

Since VEP models often depend on specific versions of Python, deep learning frameworks, GPU libraries, and compiled third-party tools, we chose run each model in a dedicated docker container [30]. In a shared environment, each new model would constrain all existing ones, and version conflicts would become more likely as the benchmark grows. Updating one model could also change the predictions of another thereby requiring tightly coupled and ever growing test-suites. Isolation allows each model to retain the environment validated by its developers, be maintained independently, and be dispatched to the hardware it requires. The benchmarking pipeline itself runs in a separate, lightweight environment and interacts with models only through the entry point and the files they write. Each model project contains the model card, the evedesign wrapper, a dependency specification, and two container definitions, a `Dockerfile` for Docker and a `Singularity.def` file for Apptainer/Singularity, which is often the only container runtime available on HPC clusters [31] (Supplementary Listing S3). This approach follows community efforts such as BioContainers, which distribute bioinformatics tools as standardized container images [32].

For the initial release, we include 12 zero-shot and 4 supervised models (Supplementary Table S2). We prioritized widely used baselines and well-established models that together cover the main classes of VEP methods. These inlcude a substitution matrix baseline, MSA-based models, protein language models, and structure-based and inverse folding models to supervised regressors ranging from linear models on one-hot encodings to Gaussian processes with composite kernels. To verify that the models behave as in the original benchmark, we compared the zero-shot Spearman correlations obtained with the PGF with those reported in ProteinGym v1.3 on the 153 datasets shared between both versions (Supplementary Table S3). ESM-2 reproduced the published correlations within rounding at all five model sizes. SaProt and RSALOR differed by a mean absolute difference of at most 0.022. We observed larger deviations for the two MSA-based Potts models, EVmutation and the site-independent model, and for ProteinMPNN, which scored higher than in ProteinGym v1.3.

### 2.4 Benchmarks run as parameterized and cached pipelines

`proteingym-benchmark` orchestrates benchmarking experiments with Data Version Control (DVC) [33], which, unlike general-purpose workflow managers (Snakemake and Nextflow) [34, 35], versions data and pipeline outputs alongside the code in git. An experiment is declared as a pipeline of stages, each defined by a command, its inputs, and its outputs. DVC links stages whose outputs are the inputs of others into a directed acyclic graph and records a hash of every input and output. Using these hashes, it skips any stage whose inputs have not changed and reuses the cached outputs. Adding a dataset, for example, only triggers the jobs for that dataset, which makes it practical to keep a growing benchmark up to date.

Experiments differ in their selection of models, datasets, and evaluation protocol. The pipelines are parameterized by a small set of variables: the datasets (each with its PGDATA file, prediction target, and split), the models (each with its container image), the CV folds, and the metrics. The pipeline expands into one job for every combination of these elements. With this design, benchmarking a new model on a subset of datasets, or evaluating all models with a different metric, only requires editing these lists (Supplementary Section A.4).

Predictions from the two settings are not directly comparable, so supervised and zero-shot models are benchmarked in separate pipelines. As both settings share the same model interface, the two pipelines consist of the same stages (Figure 2A). We first run a prediction job for every combination of model, dataset, and fold. The predictions of each job are then scored by `proteingym-base` on both the test fold and the training folds; a large gap between the two may indicate overfitting. Metrics are subsequently aggregated over folds and summarized in a single table that is read by the web catalog. The zero-shot pipeline omits iteration over folds, as each model scores every dataset once.

Users can run the benchmark locally on their own selection of models and datasets. New models can be onboarded by opening a pull request with the model implementation to the `proteingym-benchmark` repository. The pull request triggers a CI workflow that detects added or changed models, builds their container images, and runs both pipelines on a small synthetic test dataset. The workflow then posts a summary of the scores to the pull request, so contributors and maintainers see whether the model runs before it is reviewed and merged into the public benchmark (Supplementary Section A.5).

### 2.5 Statistically rigorous web catalog

The original ProteinGym website presents a leaderboard that makes it easy to identify top-scoring models. However, users who want to inspect datasets and models in detail must search for the publications or repositories that provide them. The leaderboard also does not indicate whether differences between models are meaningful, presenting a difference in average Spearman correlation of one thousandth in the same way as a difference of one tenth.

We developed a new web catalog of benchmarking artifacts to address both issues. The catalog parses the model cards and dataset manifests to describe models and datasets automatically, links to the Hugging Face repository from which datasets can be downloaded, and reads benchmarking results directly from the `proteingym-benchmark` repository, so that new results appear as soon as the benchmark is updated. Results can be filtered by molecular property; datasets can also be grouped by protein or by publication, which limits the influence of heavily studied proteins on comparisons (Figure 2B, Supplementary Figure S1). Each comparison is restricted to the datasets that every displayed model has scored.

The catalog also tests whether differences between models are significant, using critical Difference test and bootstrapped deltas (Supplementary Section A.6). The first test is rank based. Models are ranked on every dataset, and two models are considered different if their mean ranks differ by more than the Nemenyi critical difference at *α* = 0.05 [36]. This test asks whether one model wins consistently across datasets and is insensitive to differences in score scale between datasets. However, it discards the size of the differences, and its outcome for a pair of models depends on which other models are included in the comparison [37]. The second test quantifies effect size by resampling the per-dataset differences between a model and the best model with a paired bootstrap [38, 39], which yields the mean and median difference with a 95% confidence interval, reported overall and for each phenotype.

### 2.6 New evaluation protocols require only isolated extensions

The PGF simplifies the addition of new evaluation protocols. As an example, we set up an experiment for a common protein engineering scenario: selecting the model best suited to find variants with property values beyond the training distribution.

We compared three supervised models, PLS, RITA regressor, and Kermut [7], on eight randomly selected datasets with sequences shorter than 150 residues. We defined hit variants as variants whose property values exceed those of all training variants, and measured recovery as the fraction of the top-*k* measured test variants that a model also ranks among its top-*k* predictions, with *k* equal to the number of hit variants, which is stored in the metadata of each split (Figure 3A). To ensure that every test set contains hit variants, we used a quantile-based split that first divides each dataset into a lower (75%) and an upper (25%) quantile. The training set is a uniform sample of 80% of the lower quantile, and the test set consists of the remaining 20% of the lower quantile and a uniform 20% sample of the upper quantile, i.e. the hit variants.

**Figure 3:**
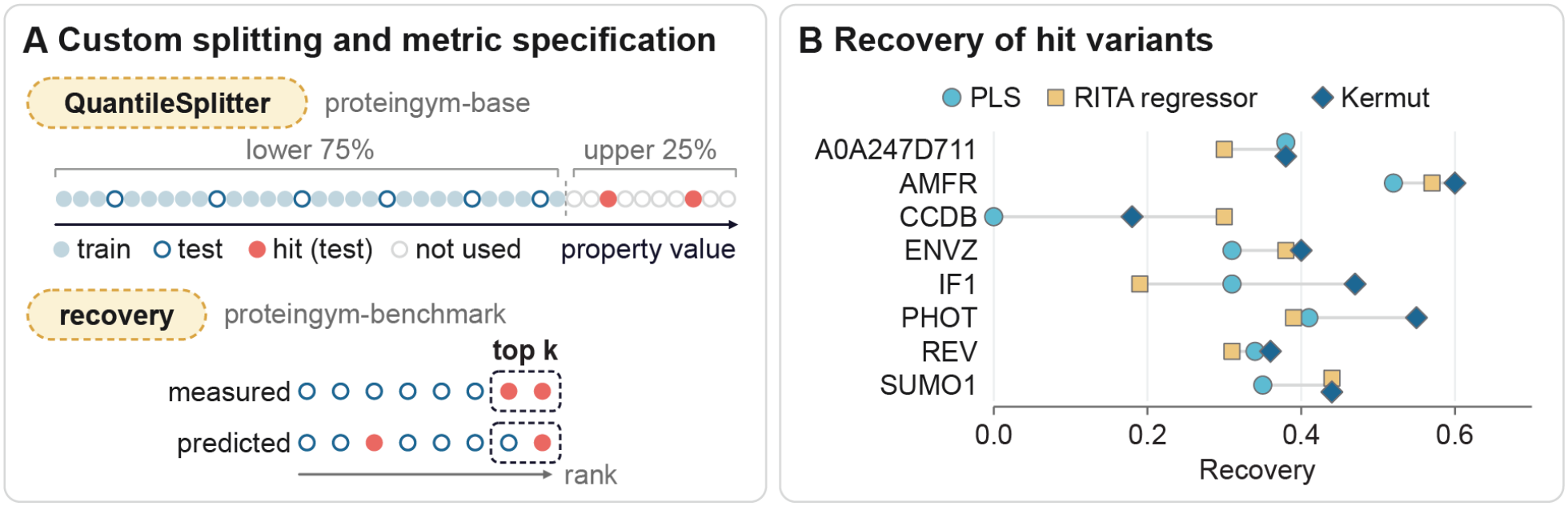
Custom evaluations. (A) Customizations for the property extrapolation experiment. The quantile splitter places all hit variants (variants exceeding the training distribution) in the test set, and the recovery metric measures the fraction of the top-*k* measured variants that a model also ranks in its top *k*. (B) Recovery of hit variants by PLS, RITA regressor, and Kermut on eight datasets (Supplementary Table S4).

The experiment required two isolated additions to `proteingym-base`, the quantile splitter and the recovery metric, together with an update of the pipeline variables. We did not have to change any model code or pipeline definitions, and both additions are now available for all models and datasets. Kermut obtained the highest or tied-highest recovery on seven of the eight datasets, with a mean recovery of 0.42, compared with 0.36 for RITA regressor and 0.33 for PLS (Figure 3B, Supplementary Table S4). These results suggest that Kermut is better suited to finding hit variants, but eight datasets are too few to rank the three models with confidence; a ranking would require the significance tests described above on a larger selection of datasets.

## 3 Discussion

We have rebuilt the framework behind ProteinGym around standardized datasets and models, parameterized benchmarking pipelines, and a web catalog with statistical model comparison. Our aim is to make it easier to compare new models with existing ones, to design new experiments, and to run meta-analyses over benchmarking results.

A central design choice of the framework is how models are standardized. Libraries such as Hugging Face Transformers reimplement every model within a single shared codebase, which yields uniform, efficient, and readily composable implementations. However, every model must then be ported and maintained by the library developers, ports may diverge from the original implementation, and models that do not fit the library’s abstractions are difficult to include. We wrap each model’s original code behind a thin interface and isolate it in a container. This approach is generally cheaper for contributors, keeps predictions faithful to the authors’ implementations, and accommodates arbitrary dependencies, at the cost of a larger compute and storage footprint per model. Because the contract fixes only the interface, the PGF could adopt other model frameworks in the future, as it has adopted evedesign.

The same design is intended to make VEP more accessible to agentic science. In other fields, benchmarks that provide standardized tasks, isolated execution environments, and automated scoring have become the main instruments through which AI agents are developed and compared [41, 42], including ARC for abstract reasoning [43], SWE-bench for software engineering [44], and Terminal-Bench, which evaluates agents on tasks in containerized environments with automated verification [45]. Progress on these benchmarks has been rapid, in part because agents can iterate against them without human intervention. Agents are also increasingly applied to scientific discovery [46, 47]. The PGF is designed to provide the same ingredients for VEP. Without the PGF, we expect an agent tasked with comparing models on a selection of datasets to spend much of its budget on reformatting data and on installing and training models. With the PGF, the agent would only have to implement the model interface to obtain a reproducible evaluation in which differences from existing models are qualified by significance tests. The experience of these fields also suggests that intensively optimized benchmarks risk saturation and overfitting [48, 42], so datasets should be added continuously, and rankings should be reported with their statistical support.

Several extensions are planned for future releases. First, we will extend the web catalog, which is at an early stage, with functionality to download and visualize predictions. Second, we will develop a taxonomy of VEP prediction tasks and use cases to organize benchmarks by the application they inform, such as clinical variant interpretation or the property extrapolation scenario described here. In protein engineering in particular, design objectives usually involve multiple properties, sometimes under variable assay conditions, which makes multi-objective prediction and support for non-sequence input features high priorities. Third, we will deepen the integration between evedesign and ProteinGym. Our goal is for models benchmarked in ProteinGym to be directly available for evedesign workflows, and for models contributed to evedesign to be benchmarked automatically. Finally, we plan to make pipeline execution environment agnostic, which would allow stages to be dispatched to HPC clusters or cloud instances. The Apptainer/Singularity definitions are a first step in this direction.

**Resources** `proteingym-base` and `proteingym-benchmark` are available at https://github.com/ProteinGym/proteingym-base and https://github.com/ProteinGym/proteingym-benchmark, respectively. The web catalog is hosted at https://proteingym.github.io/proteingym-benchmark/, and datasets are hosted at https://huggingface.co/datasets/ProteinGym/ProteinGym2.0.

## A Supplementary Materials

### A.1 Dataset schema

The `manifest.toml` file follows a hierarchical schema of required and optional components. Each top-level component is referenced by a key and defined by one of three value types, namely single values, tables, or arrays of tables (Table S1). Tables allow nested definitions of dataset components, e.g. the `publication` key can be described by a title, a DOI, and a journal. Arrays of tables allow multiple entries under the same key, so that a single dataset can contain multiple assays or structures. An example manifest is shown in Listing S1. The schema shown here corresponds to the version of `proteingym-base` described in this work; the current schema is documented online.

**Table S1:**
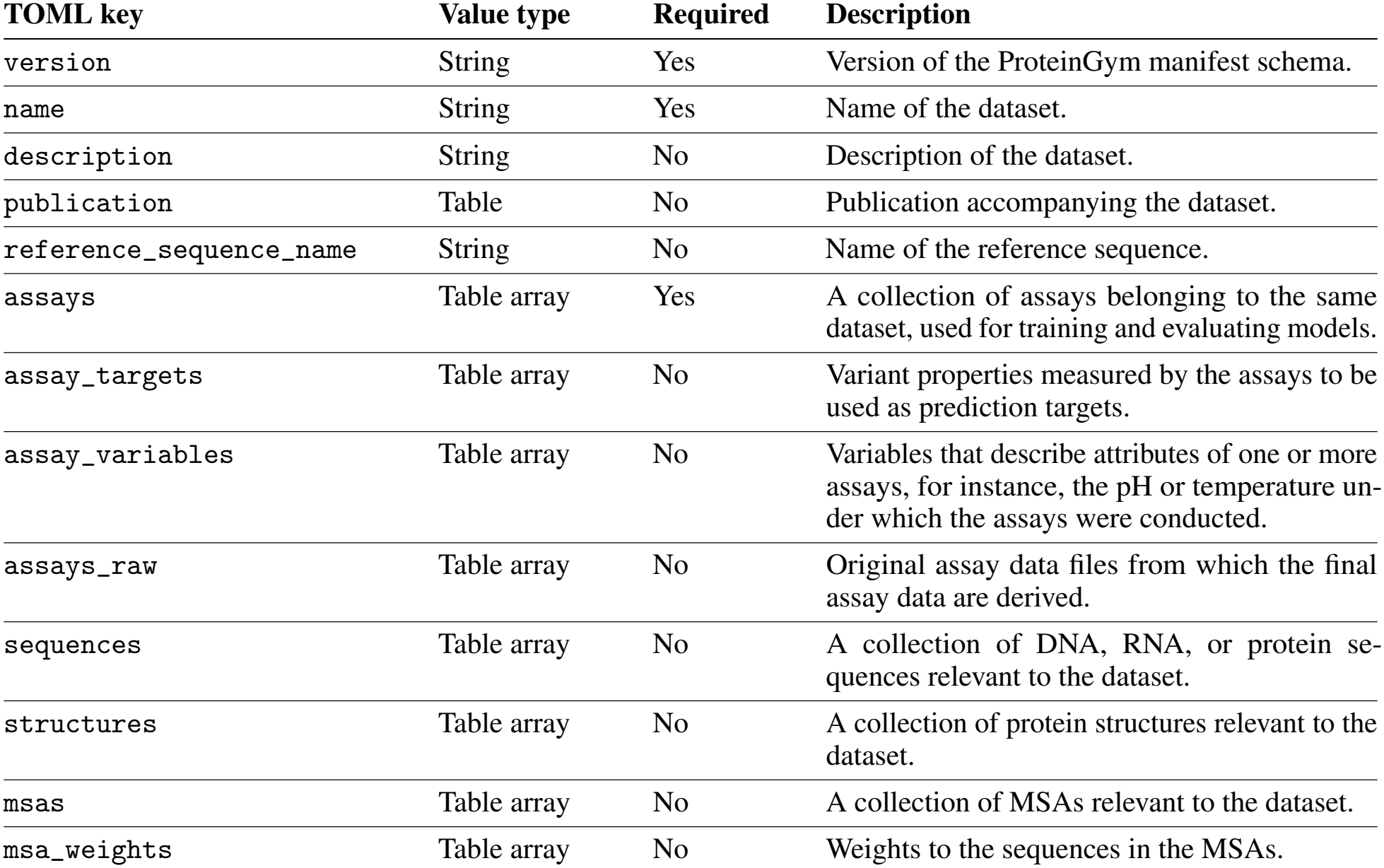
Top-level structure of the manifest.toml file.

| TOML key | Value type | Required | Description |
|---|---|---|---|
| version | String | Yes | Version of the ProteinGym manifest schema. |
| name | String | Yes | Name of the dataset. |
| description | String | No | Description of the dataset. |
| publication | Table | No | Publication accompanying the dataset. |
| reference_sequence_name | String | No | Name of the reference sequence. |
| assays | Table array | Yes | A collection of assays belonging to the same dataset, used for training and evaluating models. |
| assay_targets | Table array | No | Variant properties measured by the assays to be used as prediction targets. |
| assay_variables | Table array | No | Variables that describe attributes of one or more assays, for instance, the pH or temperature un-der which the assays were conducted. |
| assays_raw | Table array | No | Original assay data files from which the final assay data are derived. |
| sequences | Table array | No | A collection of DNA, RNA, or protein se-quences relevant to the dataset. |
| structures | Table array | No | A collection of protein structures relevant to the dataset. |
| msas | Table array | No | A collection of MSAs relevant to the dataset. |
| msa_weights | Table array | No | Weights to the sequences in the MSAs. |

**Listing S1:**
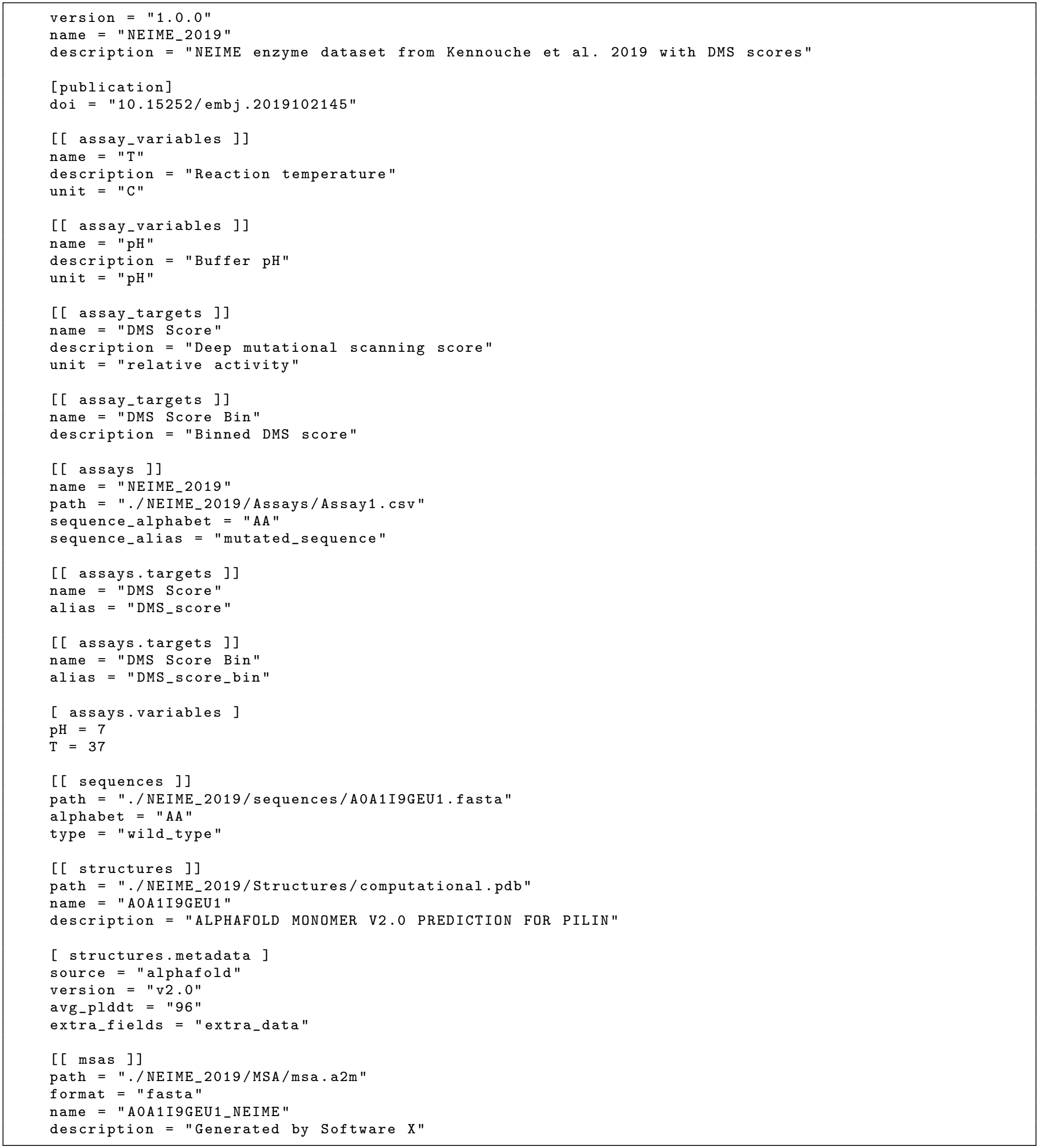
Example `manifest.toml` file for documenting and building the enzyme dataset from Kennouche et al. 2019. TOML files are structured data formats that store key-value pairs. Single bracket syntax (e.g. `[publication]`) defines tables, which are themselves collections of key-value pairs. Double brackets (e.g. `[[assay_variables]]`) define arrays of tables, allowing multiple entries to be stored under the same key. Each time `[[assay_variables]]` appears, it creates a new element in the array. Tables can be nested within array elements using dot notation. For instance, `[[assays.targets]]` nests a targets array within the most recently defined `assays` entry, while `[assays.variables]` nests a single variables table. Similarly, `[structures.metadata]` nests a metadata table under the most recently defined `structures` entry.

The PGDATA archive built from a manifest contains `manifest.lock`, the machine-generated, resolved manifest describing all metadata and the archive-internal paths of every artifact, which is not meant to be modified; `assays/`, CSV files of the assay data with at least a column of variant sequences and one or more target columns; `assays_raw/`, optional original assay files from which the assay data are derived; `sequences/`, DNA, RNA, or protein sequences in FASTA or FASTQ format; `structures/`, structures in PDB, CIF, or mmCIF format; and `msas/`, MSAs in A2M or A3M format together with sequence weights in NPY format.

### A.2 Models in the initial release

**Table S2:**
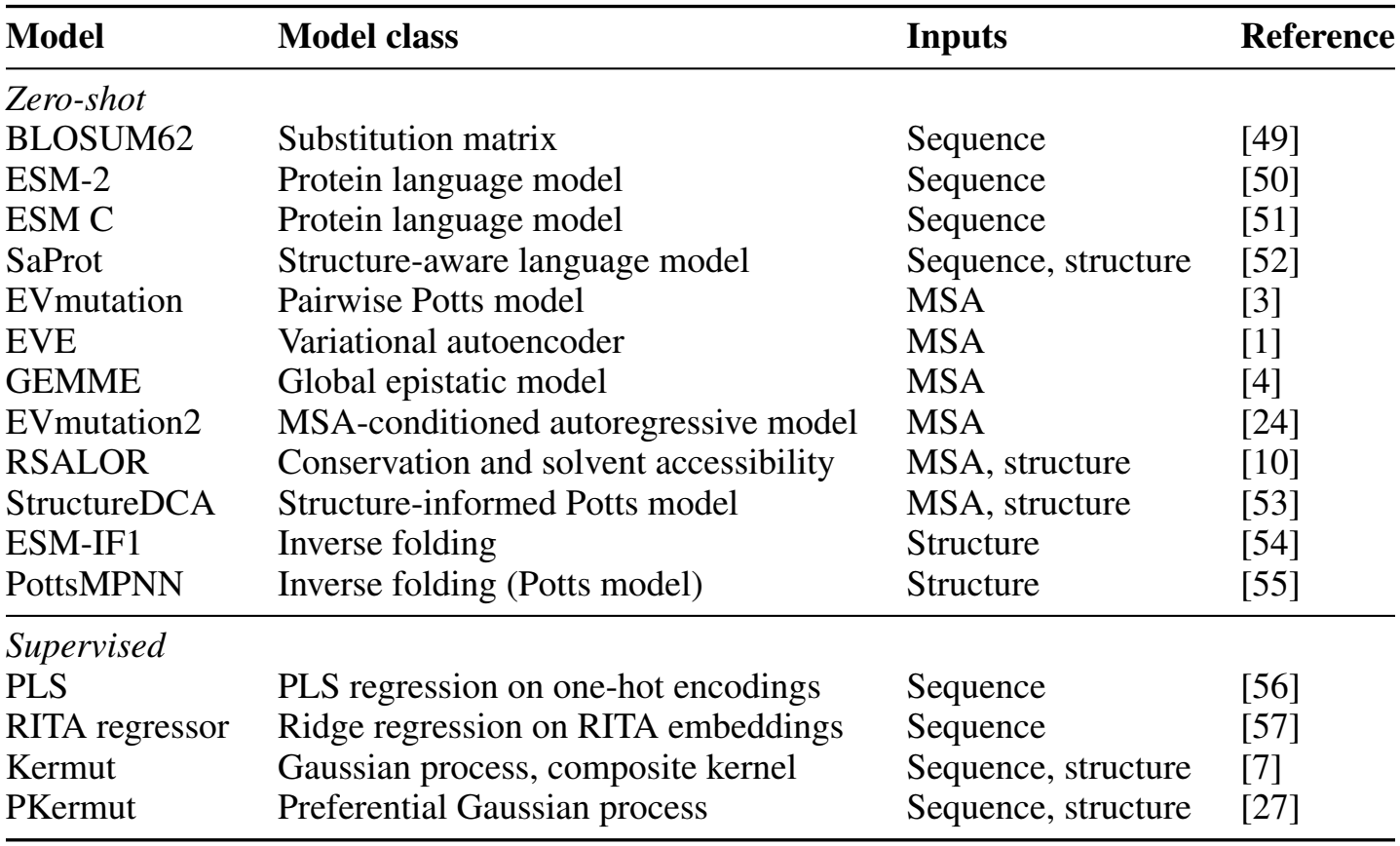
Models available in the initial release of `proteingym-benchmark`. Inputs list the data used by each model in addition to the variant sequences; supervised models additionally use the assay labels of the training folds. All models are implemented as evedesign wrappers and run through the shared entry point.

| Model | Model class | Inputs | Reference |
|---|---|---|---|
| *Zero-shot* |  |  |  |
| BLOSUM62 | Substitution matrix | Sequence | [49] |
| ESM-2 | Protein language model | Sequence | [50] |
| ESM C | Protein language model | Sequence | [51] |
| SaProt | Structure-aware language model | Sequence, structure | [52] |
| EVmutation | Pairwise Potts model | MSA | [3] |
| EVE | Variational autoencoder | MSA | [1] |
| GEMME | Global epistatic model | MSA | [4] |
| EVmutation2 | MSA-conditioned autoregressive model | MSA | [24] |
| RSALOR | Conservation and solvent accessibility | MSA, structure | [10] |
| StructureDCA | Structure-informed Potts model | MSA, structure | [53] |
| ESM-IF1 | Inverse folding | Structure | [54] |
| PottsMPNN | Inverse folding (Potts model) | Structure | [55] |
| *Supervised* |  |  |  |
| PLS | PLS regression on one-hot encodings | Sequence | [56] |
| RITA regressor | Ridge regression on RITA embeddings | Sequence | [57] |
| Kermut | Gaussian process, composite kernel | Sequence, structure | [7] |
| PKermut | Preferential Gaussian process | Sequence, structure | [27] |

**Table S3:** Comparison of zero-shot Spearman correlations obtained with the PGF and those reported in ProteinGym v1.3 [18]. Datasets were matched by identifier, which yields 153 datasets shared between both versions; the re-curated stability datasets of Tsuboyama et al. carry new identifiers in the PGF and are not included. *n* is the number of shared datasets scored by the model in both versions. Δ denotes the per-dataset difference in Spearman correlation (PGF minus ProteinGym v1.3), and *r* the Pearson correlation of per-dataset scores between both versions. ProteinGym v1.3 scores are reported to three decimals, so that differences below 0.0005 are not resolved. SaProt is compared with the variants that use AlphaFold2 structures. ProteinMPNN was replaced with LigandMPNN, leading to a 0.157 mean spearman increase from ProteinGym v1.3. EVmutation and its site-independent model were fit using mmseqs2 MSAs (as opposed to jackhmmer) and *θ* = 0.9, as opposed to the original *θ* = 0.8.

| **Model** | *n* | **Mean Spearman** |  | **Mean Δ** | **Mean \|Δ\|** | **Max \|Δ\|** | *r* |
|---|---|---|---|---|---|---|---|
|  |  | **PGF** | **v1.3** |  |  |  |  |
| Site-independent | 133 | 0.348 | 0.374 | -0.027 | 0.099 | 0.653 | 0.53 |
| EVmutation | 133 | 0.375 | 0.423 | -0.048 | 0.092 | 0.631 | 0.62 |
| ESM-2 (8M) | 134 | 0.183 | 0.183 | 0.000 | 0.000 | 0.000 | 1.00 |
| ESM-2 (35M) | 134 | 0.273 | 0.273 | 0.000 | 0.000 | 0.000 | 1.00 |
| ESM-2 (150M) | 134 | 0.358 | 0.358 | 0.000 | 0.000 | 0.000 | 1.00 |
| ESM-2 (650M) | 134 | 0.406 | 0.406 | 0.000 | 0.000 | 0.000 | 1.00 |
| ESM-2 (3B) | 134 | 0.404 | 0.404 | 0.000 | 0.000 | 0.000 | 1.00 |
| SaProt (35M) | 147 | 0.341 | 0.344 | -0.003 | 0.020 | 0.127 | 0.98 |
| SaProt (650M) | 147 | 0.418 | 0.425 | -0.007 | 0.022 | 0.269 | 0.98 |
| ProteinMPNN | 148 | 0.332 | 0.175 | +0.157 | 0.159 | 0.378 | 0.80 |
| RSALOR | 147 | 0.446 | 0.443 | +0.003 | 0.022 | 0.103 | 0.98 |

### A.3 Model contract and model projects

The shared entry point of evedesign-wrapped models exposes a `train` command, which builds and scores a model using the arguments listed below.

`model_card_file` Path to the model card (`README.md`); defaults to the model card copied into the container.

`dataset_file` Path to a PGDATA dataset, including its splits.

`target` Name of the assay target to train and evaluate on.

`split` Name of the splitting strategy; omitted for zero-shot models.

`test_fold` Index of the fold used as the test set; omitted for zero-shot models.

The entry point reads the model card, resolves the evedesign class named in its `model_class` field, and instantiates it with the hyperparameters in the model card. Models tagged `zero-shot` are built without training data and score the full dataset. Supervised models are built on the training folds and score both the test and the training folds, so that metrics can also be computed on the training data. Predictions are written as a PGDATA prediction file to the container’s output directory. An example model card is shown in Listing S2, and the structure of a model project in Listing S3.

**Listing S2:**
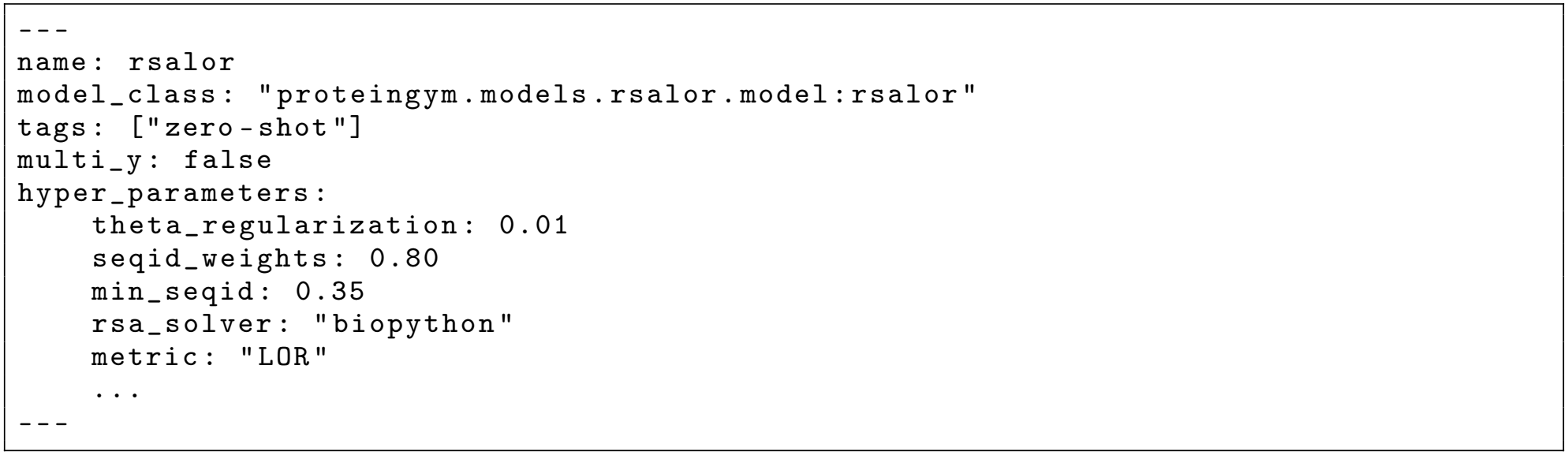
Front matter of the model card of the RSALOR model [10] (abridged). The `model_class` field names the evedesign wrapper class that the shared entry point instantiates with the given hyperparameters, and the `tags` field declares the prediction setting. The Markdown body that follows the front matter documents the model in human-readable form.

**Listing S3:**
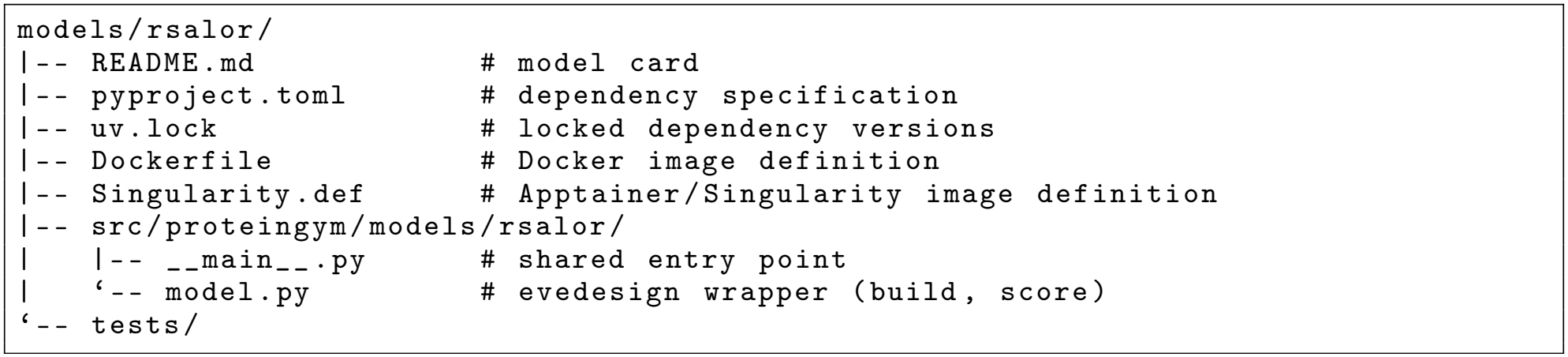
Structure of a model project in the `models` directory of `proteingym-benchmark`. The model card doubles as the model’s documentation; `model.py` implements the evedesign wrapper; `main .py` is the shared entry point; `pyproject.toml` and `uv.lock` pin the model’s dependencies; and the `Dockerfile` and `Singularity.def` files define equivalent container images for Docker and Apptainer/Singularity, respectively.

### A.4 Pipeline parameterization and stages

Benchmarking pipelines are parameterized by the variables listed below.

`datasets` An array of datasets stored in a `datasets.json` file (Listing S4). Each dataset has a `name`, an `input_filename` pointing to a PGDATA file, a `target` indicating the assay target to predict, and a `split` specifying the CV scheme.

`models` An array of models stored in a `models.json` file (Listing S4). Each model has a `name` and an `image` specifying its container image.

`output` Paths where predictions and metrics are stored.

`metrics` An array of metrics to score models by.

`folds` An array of fold indices of the CV scheme. Each fold index identifies the variants that make up the test set.

`score_modes` The partitions on which metrics are computed, i.e. the test fold, the aggregate of training folds, and each fold individually.

Both the supervised and the zero-shot pipeline consist of the four stages described below.

`create_training_job` Creates a matrix of all combinations of models, datasets, and folds (without folds for zero-shot pipelines), and runs the entry point of the corresponding model container for each combination.

`calculate_metric` Scores the predictions of each job with the selected metrics using `proteingym-base`, producing a JSON file of metrics per job.

`aggregate_metrics` Computes the mean and standard deviation of each metric over folds for every combination of model and dataset.

`generate_metrics_csv` Summarizes the aggregated metrics of all combinations in a single CSV file, which is read by the web catalog.

Experiments are executed with `dvc repro`, as described in the online documentation.

**Listing S4:**
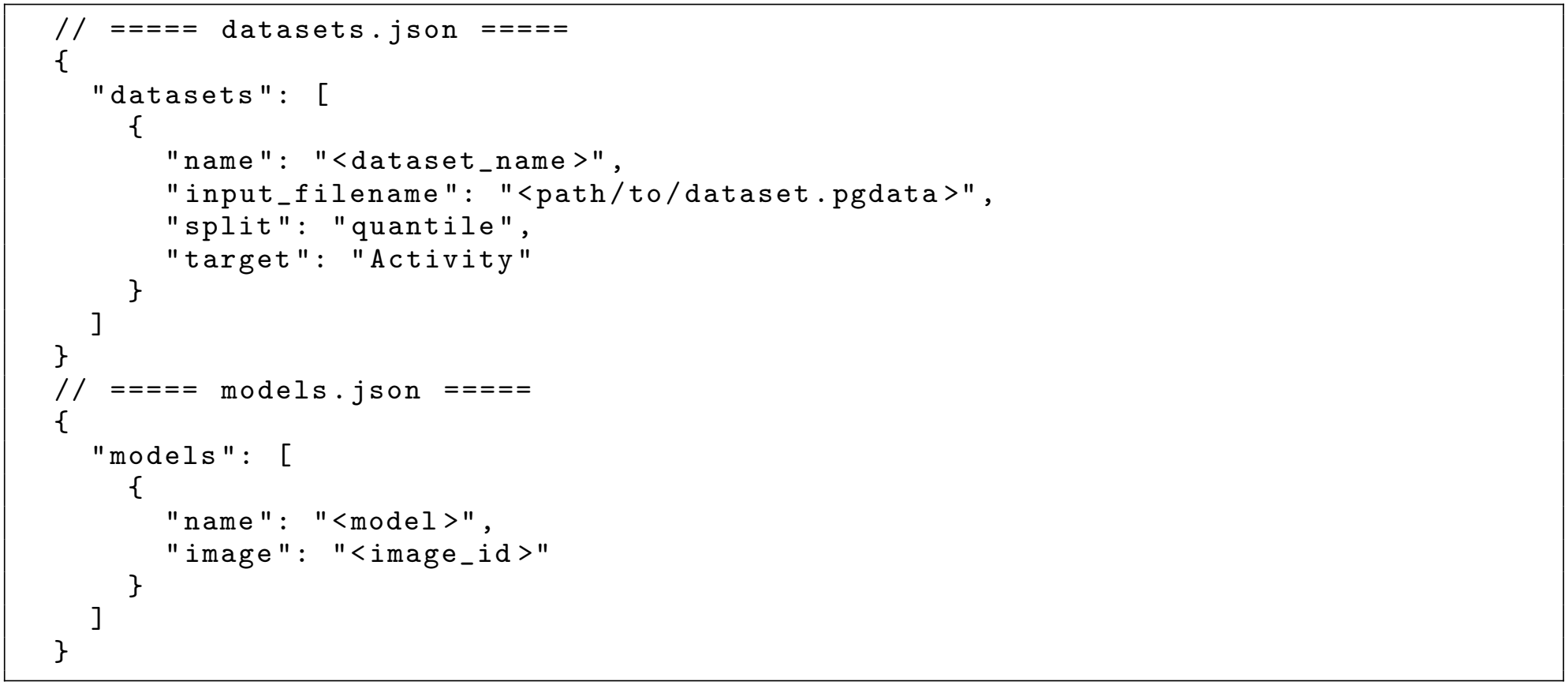
Example `datasets.json` and `models.json` files describing which datasets and models should be benchmarked. While the examples contain a single model and dataset, the `datasets` and `models` keys store arrays that may contain any number of entries. Each file is a separate JSON document; the comment headers indicate the filename and are not part of the JSON content.

### A.5 Continuous integration workflow

Pull requests to `proteingym-benchmark` trigger a GitHub Actions workflow consisting of the steps described below.

#### Install requirements

Installs the requirements of `proteingym-benchmark`.

#### Check for relevant changes

Determines which models in the `models` directory were added or changed.

#### Build or reuse model images

Builds container images for added or changed models, tagged by a hash of the model project, and pushes them to the GitHub Container Registry; images of unchanged models are reused.

#### Set up CI datasets

Copies the test dataset into the runner file system. The test dataset is a small synthetic dataset whose target is a simple function of sequence, which keeps fitting times short.

#### Run benchmarking

Runs the supervised and zero-shot pipelines for the changed models, ensuring that they complete without errors.

#### Create PR report

Posts the scores of each model, averaged across datasets, to the pull request as a sanity check.

### A.6 Statistical model comparison

Let *k* models be evaluated on *N* datasets (or groups of datasets), and let *s*_*ij*_ denote the score of model *j* on dataset *i*.

#### Nemenyi critical difference

For each dataset *i*, models are ranked by *s*_*ij*_ (rank 1 being best, ties receiving the average rank), and the mean rank 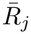 of each model is computed over datasets. Two models are considered significantly different at level *α* if their mean ranks differ by more than the critical difference

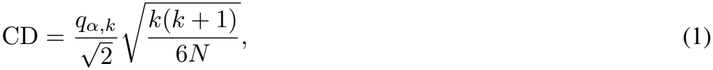

where *q*_*α,k*_ is the upper *α* quantile of the studentized range distribution for *k* groups and infinite degrees of freedom [36]. The web catalog uses *α* = 0.05.

#### Paired-delta bootstrap

For two models *a* and *b*, the per-dataset differences *d*_*i*_ = *s*_*ia*_ *−s*_*ib*_ are computed, and their mean 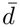 is reported with a 95% percentile bootstrap confidence interval obtained from 10,000 resamples of the *N* differences with replacement. Because each dataset contributes the scores of both models to the same resample, the test accounts for the pairing of scores. The difference is considered significant if the interval excludes zero. In the web catalog, each model is compared with the best-scoring displayed model, overall and for each phenotype. These intervals are not corrected for multiple comparisons, and because the reference is the best observed model, individual intervals that exclude zero should be interpreted with caution when many models and phenotypes are compared.

#### Grouping

Optionally, datasets measured on the same protein or reported in the same publication are first averaged into a single unit, so that *N* counts proteins or publications rather than individual datasets.

### A.7 Property extrapolation experiment

**Table S4:** Recovery scores (higher is better) of PLS, RITA regressor, and Kermut across eight ProteinGym datasets. The best score per dataset is shown in bold.

| **Dataset** | **PLS** | **RITA regressor** | **Kermut** |
|---|---|---|---|
| A0A247D711_LISMN_Stadelmann_2021 | **0.38** | 0.30 | **0.38** |
| AMFR_HUMAN_Tsuboyama_2023_4G3O | 0.52 | 0.57 | **0.60** |
| CCDB_ECOLI_Adkar_2012 | 0.00 | **0.30** | 0.18 |
| ENVZ_ECOLI_Ghose_2023 | 0.31 | 0.38 | **0.40** |
| IF1_ECOLI_Kelsic_2016 | 0.31 | 0.19 | **0.47** |
| PHOT_CHLRE_Chen_2023 | 0.41 | 0.39 | **0.55** |
| REV_HV1H2_Fernandes_2016 | 0.34 | 0.31 | **0.36** |
| SUMO1_HUMAN_Weile_2017 | 0.35 | **0.44** | **0.44** |

**Figure S1:**
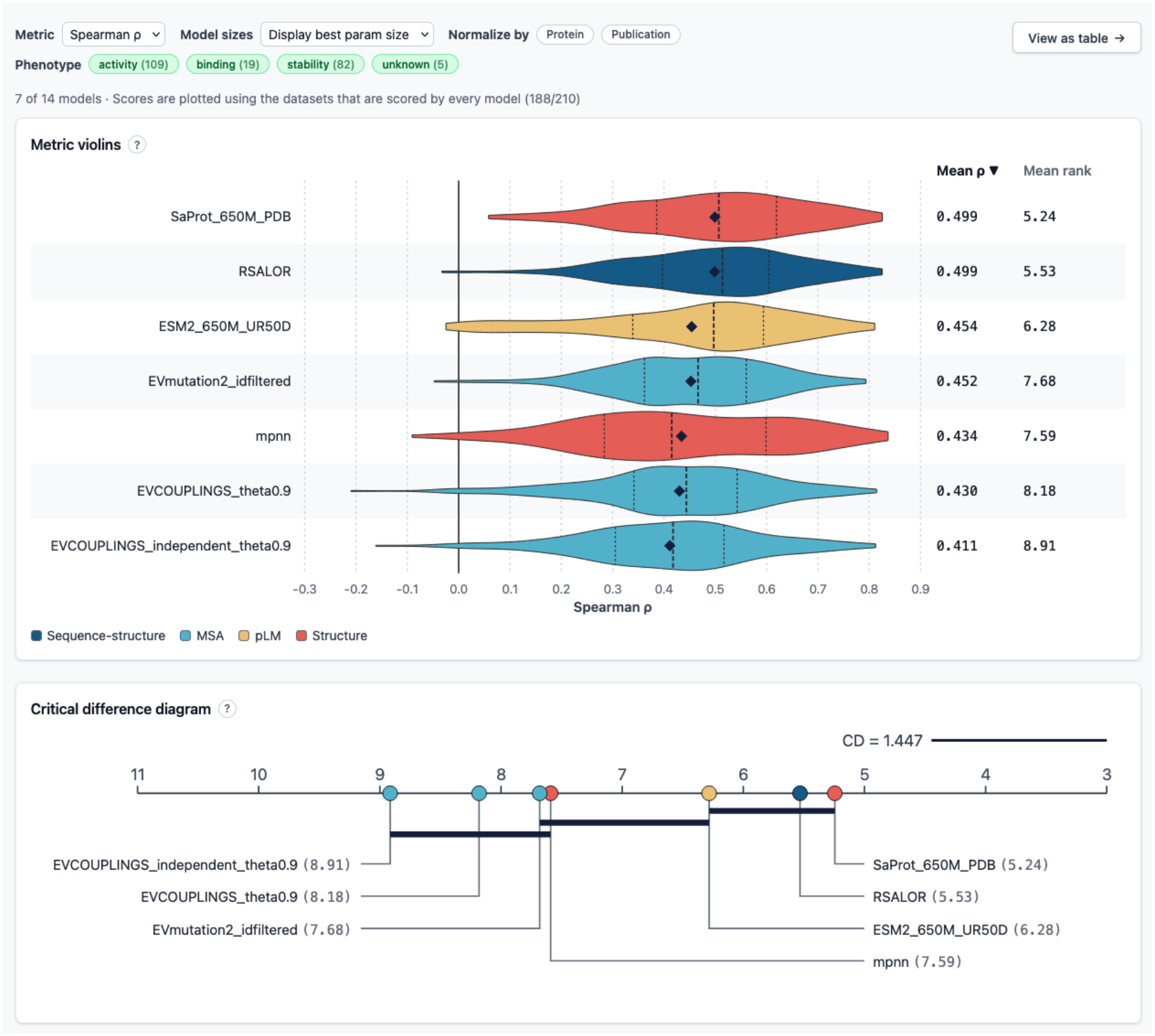
Zero-shot benchmark view of the web catalog. Results can be filtered by phenotype and model size, and datasets can be grouped by protein or publication. Top, distribution of Spearman correlations across all datasets scored by every displayed model, with the mean score and mean rank of each model. Bottom, critical difference diagram, in which models connected by a horizontal bar do not differ in mean rank by more than the Nemenyi critical difference (*α* = 0.05). All results were generated with the PGF benchmarking pipeline; some of the displayed models and model variants have not yet been added to the public model repository (Supplementary Table S2).

